# An efficient and robust HPLC method to determine the sialylation levels of human epithelial cells

**DOI:** 10.1101/2021.08.26.457765

**Authors:** Hyo Jeong Kim, Stephanie Schweiker, Katie Powell, Stephan M Levonis

## Abstract

Sialyltransferase, an enzyme responsible for attaching sialic acid to the cell surface, is reported to play a key role in cancer, making sialyltransferase a potent therapeutic target in drug development for cancer. Hence, this paper aimed to develop a simple method to detect and quantify sialic acids in cancer cells. An efficient method was developed using a reverse-phase ion-pairing HPLC-UV using triisopropanolamine as the ion-pairing agent with a C18 column. Neu5Ac was successfully eluted with the retention time 6.344 min with a flow rate of 0.4 mL/min. The proposed method was validated appropriately according to the AOAC guidelines (2013). This work demonstrates that the proposed method is not only relatively simple but also cost and time effective compared to pre-existing methods to successfully determine both free and protein-bound Neu5Ac in a complex cancer cell matrix. Furthermore, by applying the proposed method, a statistically significant decrease was observed for both HeLa and HuCCT1 cell lines with the application of deoxycholic acid – a known sialyltransferase inhibitor. Hence, the proposed method may be applicable to evaluate the effectiveness of sialyltransferase inhibitors.

## Introduction

Inflammation plays key roles at all stages of cancer [1]. It is the driver of cancer progression, mediating cancer cells to invade into the secondary tissues and increasing tumour survival and progression by modulating the immune responses [2]. Tumours can avoid the immune cell responses directed at them by co-opting inhibitory receptors, which leads to unresponsive T-cells [3]. The interaction between immune activation and cancer pathogenesis have been investigated as a target for potential immunotherapy [4]. Hence, in recent years, inhibitors with the ability to break the cycle of immune suppression have been widely investigated in the hope of enabling the suppressed immune cells to thereby selectively target cancer cells [5].

Sialic acid (Sia) is a generic term for nitrogen or oxygen substituted derivatives of neuraminic acid – a nine-carbon monosaccharide [6]. They appear as either free or bound to protein in nature and are widely expressed as the outer terminal units of glycans on the surfaces of all vertebrate cells [7, 8]. N-acetylneuraminic acid (Neu5Ac) is the most predominant type of sia on mammalian cells, synthesised via various biological pathways in human [6]. Due to their unique location and abundance on the cell surface, sias participate in a wide variety of human pathophysiological processes, including cell-to-cell adhesion, tumour cell metastasis, and inflammation. Therefore, a robust measure of inflammatory status may be provided by measuring sia level. Additionally, there have been an increase in the number of reported cases where patients with various forms of cancer have increased sia levels. Hence, hypersialylation is one of the current emerging mechanisms under investigation as a potential anti-metastatic approach targeting inflammation [5].

Various methods and techniques ranging from colorimetry, fluorimetry to enzymatic and chromatographic analysis have been reported to determine sias [8]. These approaches hold limitations of requiring entire tissue samples to extract, lack classification amongst diverse cell types, and overlook major glycans on minor cell types. Furthermore, under normal circumstances, sias are typically found as part of a glycoconjugate (approximately 73%) rather than as a free acid [7]. Thus, for accurate analysis, sias need to be purified and completely released from the respective glycoproteins, glycolipids and oligosaccharides prior to the analysis.

An acid hydrolysis method is a popular approach to determine sias as it releases sias without causing changes in their modification from the biological sources [8]. Several published studies have used sulfuric acid to release sias but other acids such as acetic acid, trifluoroacetic acid and hydrochloric acid, are also commonly used for their availability, cost, and most importantly for their volatility; they are easily removable by lyophilization [9, 10]. Following acid hydrolysis, numerous spectroscopic approaches exist to determine sias. These approaches hold a major limitation of overestimating the concentration of sias via interferences [11]. Therefore, when developing a chromatographic method, there is a substantial focus on separating sias from the interfering compounds. One proven approach that is sensitive, specific and applicable to most sias is via derivatization with 1,2-diamino-4,5-methylenedioxybenzene dihydrochloride (DMB) followed by a high-performance liquid chromatography (HPLC) analysis with fluorescent detection [12]. However, despite an improvement in the chromatographic separation via DMB-labeled sias, the derivatization methods require reagents that are costly and often not time efficient.

Hence, despite advancements, there are many challenges currently associated with determining sias in a complex matrix, such as in cancerous cells. This research will outline a reverse-phase ion pair HPLC method for exploring these limitations via simple precipitation of sias with ethanol followed by acid hydrolysis to isolate the sias bound to protein. The method will then be evaluated by measuring changes in sia level in various cancerous cell lines with and without known inhibitor.

## Materials and Methods

### Chemicals, reagents, and materials

The chemicals: Triisopropanolamine (TIP); phosphoric acid; hydrochloric acid (HCl); sodium hydroxide (NaOH) and ethanol (HPLC grade) were purchased from Sigma-Aldrich (Australia). The standard: Neu5Ac (HPLC grade, ≥97%) and Neu5Gc (HPLC grade, ≥95%) were also purchased from Sigma-Aldrich (Australia). Filter paper (0.45 μm, 25 mm Diameter Hydrophobic PTFE) was manufactured from Aijiren Corp (Quzhou, China). All standard solutions and the mobile phase (TIP solution) were prepared with deionized water obtained from Milli-Q water system (Millipore S.A.S 67120 Molsheim France).

For cell culture, Fetal Bovine Serum (FBS), Dulbecco’s Modified Eagle Medium (DMEM), Streptomycin/Penicillin and TrypLE were purchased from Gibco by life technologies (Denmark). Pre-made Phosphate buffer solution (PBS) at appropriate concentrations and pH was purchased Sigma-Aldrich (Australia).

### Instrumentation

The centrifuge was used widely across the preparatory step. A pH probe was used to buffer the TIP solution accordingly. Calibrated automatic pipettes and analytical balance were used for the volume and weight measurement. The mass of the cells was measured by a XSE105 Dual Range Mettler Toledo weighing balance.

The samples were detected and quantified via a reverse phase HPLC. All HPLC analyses were performed on a HPLC Waters Alliance e2695 Separation Module (Waters Corporation, Milford, MA, USA) coupled to a Waters 2489 UV/Visible Detector set at 215 nm. The separation was performed on the C_18_ column Luna 5 μm (2) 100 A 250 x 4.5 mm (Phenomenex Inc., Torrance, CA, USA).

### Chromatographic condition of HPLC

All separations were performed isocratically at 25°C with a mobile phase comprised of 50 mM TIP solution buffered to pH 3.5 via phosphoric acid (99% Sigma-Aldrich, Australia). The flow rate was maintained at 0.4 mL/min. A 10 μL sample volume was injected directly for all experiments with two repeated injection of a 10 μL blank TIP buffer solutions between each sample runs to reduce possible interference peaks. The sias eluted from the column were monitored by a UV/VIS spectrophotometer set at 215 nm. Peak identification was performed by comparing the retention times and absorption spectra of the respective standard solutions with the prepared samples.

### Preparation of standard and stock solution

The standard stock solutions of Neu5Ac (1 mM) and Neu5Gc (0.5 mM) were prepared by the addition of 30.9 mg of Neu5Ac and 16.3 mg of Neu5Gc in 100 mL flask. TIP buffer solution (50 mM) was added to reach a final volume of 100 mL and stored at 4°C in a fridge. The working standard solutions were produced by diluting the respective stock solution in 50 mM TIP buffer solution to form 0.03125, 0.0625, 0.125, 0.25, 0.5 and 1 mM for Neu5Ac and 0.03125, 0.0625, 0.125, 0.25 and 0.5 mM for Neu5Gc. The working standard solutions were freshly prepared before the start of the analysis via serial dilution using the stock solution.

### Sample collection and preparations

#### Culturing conditions for the cells

Total of two carcinoma cell lines: human epithelial cervical adenocarcinoma (HeLa; American Type Culture Collection) and the human liver bile duct carcinoma (HuCCT1; Cellbank Australia) were used for the experiment. The HeLa cell line was supplied by the ATCC (Manassas, USA) as catalogue number ATCC^®^ CCL-2^™^, and was purchased from In Vitro Technologies (Noble Park North, VIC, Australia). The HeLa cell line has been described before [13]. The HuCCT1 cell line was supplied by the JCRB Cell Bank (Osaka, Japan) as catalogue number JCRB0425, and was purchased from CellBank Australia (Westmead, NSW, Australia). The HuCCT1 cell line has been described before [14].

The culturing conditions for both HeLa and HuCCT1 cell lines were optimised. The cells were cultured in DMEM supplemented with 10% FBS and 0.1% Streptomycin/Penicillin in T-75 flask. The cells were maintained at 37°C in a humidified atmosphere of 5% carbon dioxide. Cells were prepared at 90 to 100% confluency. Once cells have reached its full confluency, cells were incubated for 15 minutes with TrypLE to be detached from the flask. Cells were washed with PBS solution and centrifuged prior to the sample preparation. The cells were kept in a freezer at negative 80°C until analysis. Cells were thawed to room temperature (25°C) prior to the analysis.

### Cell preparations

#### Free sialic acids

Prior to the sample preparation step, the cultured cells were washed with 1 mL of phosphate-buffered saline (PBS) solution in attempt to remove media. Ethanol (2 mL) was added to the washed samples to precipitate free sias. The solution was inverted (4 times), vortexed (1 min) and rested at room temperature (25°C) for 10 minutes. It was then centrifuged (150 g, 25°C) for 5 minutes. The supernatant containing the free sias was dried completely using a rotary evaporator under high vacuum to eliminate ethanol contamination. The mass of the cells was obtained by measuring the vacuum reduced supernatant. The 0.3 mL of TIP buffer solution was added to the reduced supernatant. The solution was filtered using a filter paper and a syringe before the HPLC injection.

#### Protein-bound sialic acids

The remaining pellet from the first centrifuge was hydrolysed to release protein-bound sias. The pellet was crushed and mixed with 1 mL of HCl (120 mM). It was heated for an hour at 80°C. After the hydrolysis step, the solution was neutralized to a pH of 7.0 with 1 mL of NaOH (120mM). Ethanol (4 mL) was added to the mixture before it was inverted (4 times), vortexed (1 min) and rested at room temperature (25°C) for 10 minutes. The hydrolysed solution was centrifuged (150 g, 25°C) for 5 minutes. The supernatant was reduced completely using a rotary evaporator under high vacuum to remove ethanol. Mass of the cell was measured. The 1 mL of TIP buffer solution was added to the vacuum reduced supernatant prior to the HPLC analysis. The solution was filtered before the HPLC injection step.

### Method Validation

The performance of the HPLC-UV method to determine sias in cultured cells was validated in terms of linearity, selectivity, accuracy (recovery), repeatability precision, Limit of Detection (LOD) and Limit of Quantification (LOQ) according to the 2013 Association of Official Analytical Chemists (AOAC) international guideline.

#### Linearity

Linearity was calculated by three injections of different Neu5Ac (0.03125, 0.0625, 0.125, 0.25, 0.5 and 1 mM) and Neu5Gc (0.03125, 0.0625, 0.125, 0.25 and 0.5 mM) concentrations where the average peak areas were plotted against concentrations. Coefficient of correlation (R^2^) obtained from the respective calibration curve was used to evaluate the linearity. According to the AOAC guideline, the acceptable value for R^2^ is greater than 0.998.

#### Selectivity

Selectivity was determined by examining the chromatograms to verify the absence of interfering peaks in the prepared samples.

#### Accuracy (recovery)

The accuracy of the method was evaluated by the percent recovery (R%) of the spiked analyte. The standard stock solution (0.5 mM) was spiked (10, 20, 30, 40 and 50 μL) five times individually into the three prepared samples with low, medium and high concentration of sias. The spiked samples were injected in triplicate.

The percent recovery of the analyte was calculated using the formula:

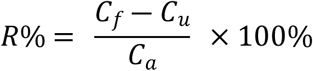

where *C_f_* represents the amount of sias quantified after the addition of the standard stock solution, *C_u_* represents the amount of sias quantified prior to spiking and *C_a_* refers to the amount of standard stock solution added. Acceptable recovery according to the guideline is within 80 to 120 % of the mean.

#### Precision

Precision was evaluated by intra-day and inter-day testings. For repeatability precision (intra-day), three replicates of the quality control samples containing all the analytes at three different concentrations (low, medium and high) were analysed simultaneously at six different times on the same day. For intermediate precision (inter-day), the analysis was performed over three successive days.

Precision was expressed as the relative standard derivation (RSD%) and was calculated using the formula:

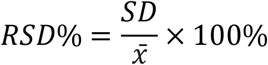

where *SD* represents the standard deviation and 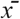 refers to the sample mean. RSD% value less than or equal to 2% was accepted according to the guideline.

#### Limit of Detection and Limit of Quantification

The LOD and LOQ was calculated using the formula:

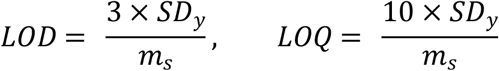

where *SD_y_* represents the standard deviation of the y-intercepts and *m_s_* refers to the average gradients from respective standard calibration curve.

### Statistical Analysis

Repeated Measures Analysis of Variance (RM ANOVA) was used to statistically compare the difference in sia level in cultured cells with/without the addition of known sialyltransferase inhibitors – deoxycholic acid. The sia levels from the control group (no addition of sialyltransferase inhibitor) were individually compared to the other conditions with varying concentration of deoxycholic acid. All statistical calculation was performed using the Statistical Package for the Social Sciences (SPSS) program. P value lower than 0.05 would indicate that the sia level in the compared cells with sialyltransferase inhibitor applied are statistically different from the control group.

## Results

### HPLC method development and optimisation

The sias in cells were detected and quantified via a reverse phase HPLC with TIP buffer solution as the ion-pairing reagent. The method was inspired from the novel study by Spyridaki and Siskos in 1999 and optimised from the results gained from the in-house research group (Levonis et al). Parameters such as chromatography conditions, mobile phase, hydrolysis conditions and sample amounts were modified to improve the analytical performance. The optimized method resulted in successful separation where Neu5Ac was eluted with a retention time of 6.344 minutes with a flow rate of 0.4 mL/min.

### Method validation

The developed method was validated as per the criteria set by the AOAC international guideline (2013).

#### Linearity

The linearity of the method was examined by the calibration curve plotted between the area of the respective peaks against the five standard concentrations of Neu5Ac over the range of 0.03125 to 1 mM (Figure 1).

**Figure 1.**
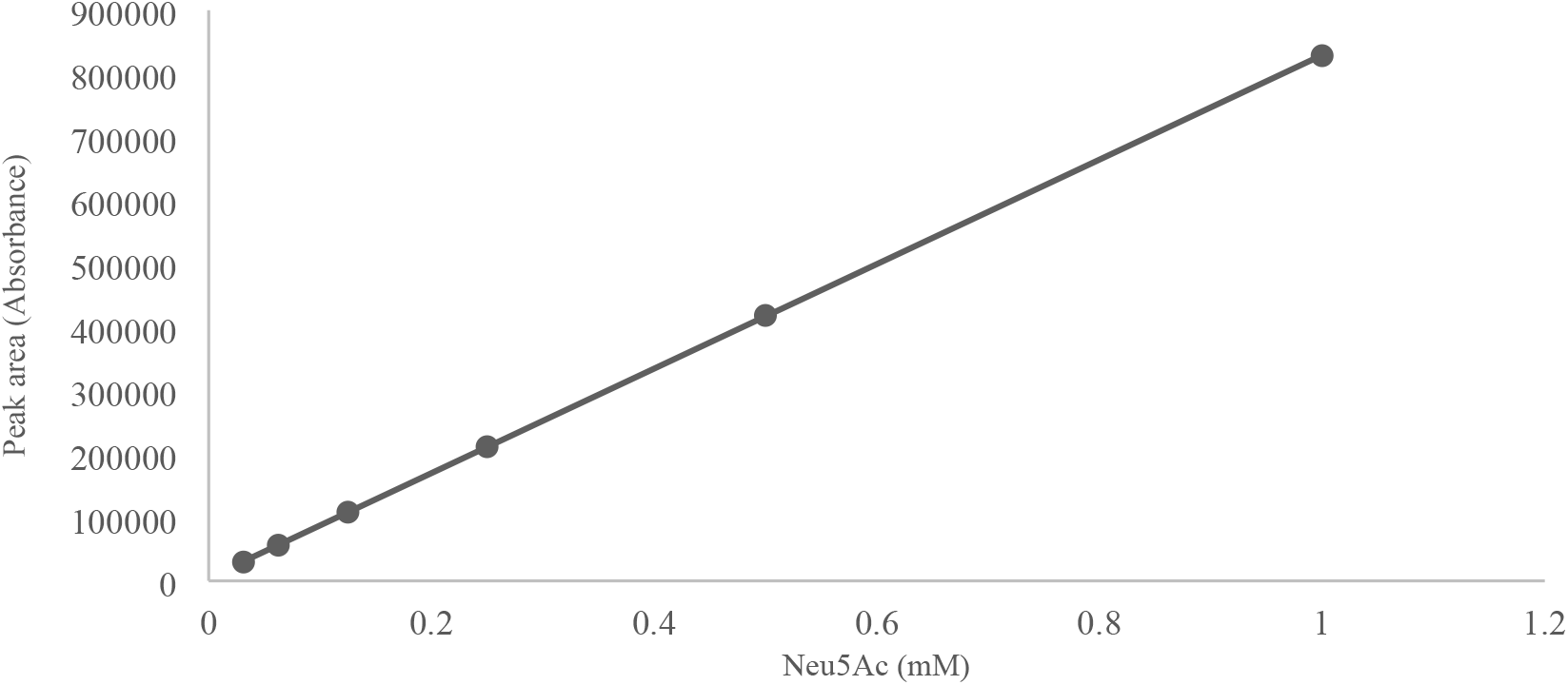
Calibration curve showing the linearity of the method: Neu5Gc.

The average R^2^ was determined to be 0.9998 for Neu5Ac (Table 1). Thus, as the R^2^ for individual analytes was achieved to be >0.998, the proposed method is linear and acceptable as it satisfies the AOAC guideline.

**Table 1.** Summary of the correlation coefficient, accuracy, precision, LOD and LOQ values for the proposed method.

|  | Neu5Ac |
| --- | --- |
| Co-efficient of Determination ($R^2 \pm SD$) | 0.9999 |
| Average accuracy (R%; n=5) | 116% |
| Average inter-day precision (RSD%, n=3) | 1.99% |
| Average intra-day precision (RSD%, n=3) | 0.85% |
| LOD (mM) | 0.002487 |
| LOQ (mM) | 0.007537 |

#### Selectivity

The proposed method was proven to have selectivity due to absence of significant interference peaks at the retention times of the Neu5Ac (Figure 2). Furthermore, by blanking with TIP solutions twice for each sample run, no carry-over effect was observed throughout the analysis.

**Figure 2.**
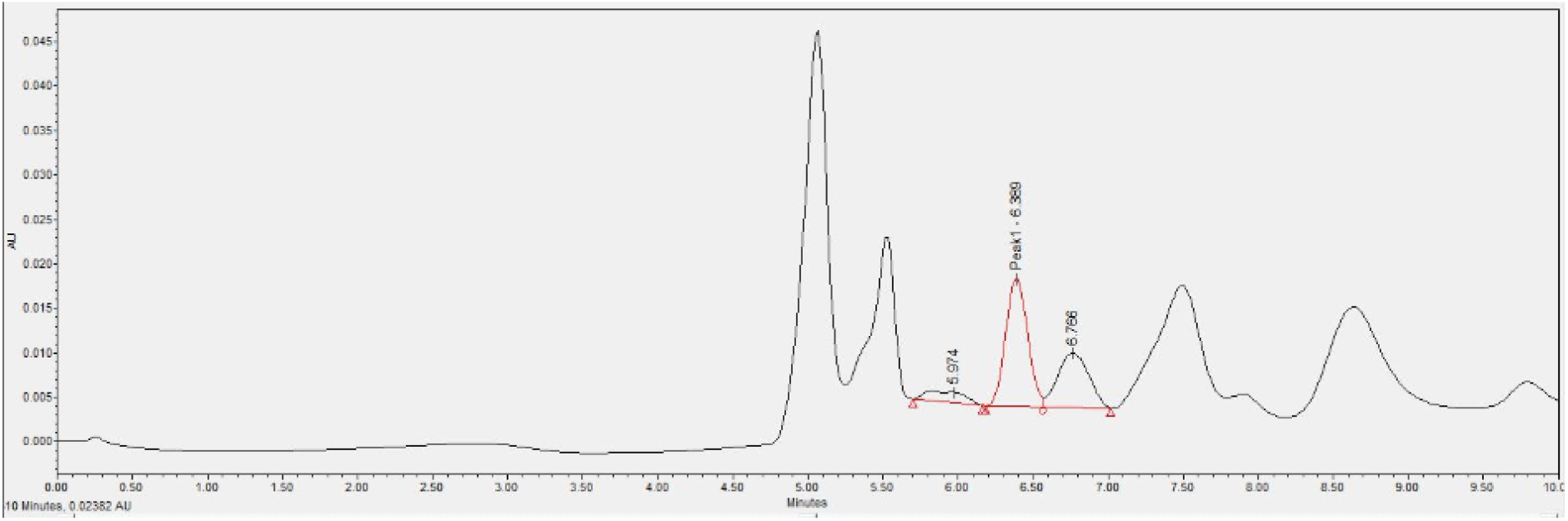
A chromatogram of cultured HeLa cells via using the optimised HPLC method. Absence of significant interference peaks at the retention times (6.344 minutes with a flow rate of 0.4 mL/min.) of Neu5Ac has been highlighted in red.

#### Accuracy (recovery)

The average recovery for high concentration of Neu5Ac sample was found to be 109%. All recovery values obtained were within the range 80 to 120%, indicating high accuracy of the proposed method.

**Table 2.**
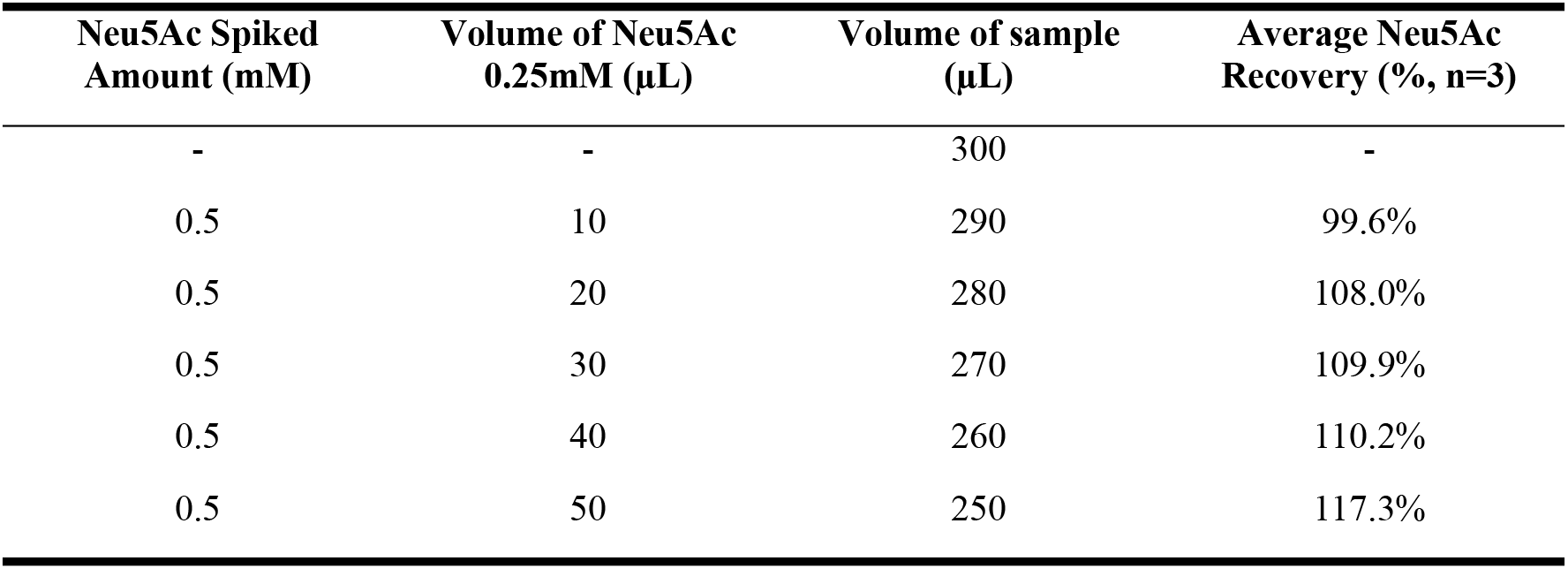
Results for accuracy (n=3) for the proposed method to determine Neu5Ac level.

| Neu5Ac Spiked Amount (mM) | Volume of Neu5Ac 0.25mM (μL) | Volume of sample (μL) | Average Neu5Ac Recovery (% , n=3) |
| --- | --- | --- | --- |
| - | - | 300 | - |
| 0.5 | 10 | 290 | 99.6% |
| 0.5 | 20 | 280 | 108.0% |
| 0.5 | 30 | 270 | 109.9% |
| 0.5 | 40 | 260 | 110.2% |
| 0.5 | 50 | 250 | 117.3% |

#### Precision

**Table 3.** Results of precision for the proposed method to determine Neu5Ac level.

| Concentration | Repeatability (Intra-day) |  |  | Intermediate precision (Inter-day) |  |  |
| --- | --- | --- | --- | --- | --- | --- |
|  | Mean (mM) | SD | RSD% | Mean (mM) | SD | RSD% |
| Low | 0.0234 | 0.00054 | 1.34% | 0.001 | 0.00000 | 0.00% |
| Medium | 0.1448 | 0.00238 | 1.65% | 0.236 | 0.00400 | 1.69% |
| High | 0.4313 | 0.00857 | 1.99% | 0.603 | 0.00513 | 0.85% |

#### LOD and LOQ

The LOD and LOQ determined for Neu5Ac were 0.002487 mM, 0.007537 mM respectively. The low LOD and LOQ of Neu5Ac enables the detection and quantification of Neu5Ac in cultured cells at low concentrations.

#### Application

By measuring sia levels in cancer cells, the effectiveness of potent inhibitors for hypersialylation can be evaluated. Hence, the developed method was applied to compare the difference in the sia level in cultured malignant cells against the cultured malignant cells with the presence of known sialyltransferase inhibitor - deoxycholic acid, for 48 hours. The HeLa and HuCCT1 cell lines were used for the application.

To determine the cytotoxicity of deoxycholic acid, malignant cells with various concentration of known sialyltransferase inhibitors (ranging from 3 to 300 μM) were dissolved in culture media and cultured for 48 hours. Preliminary qualitative results were taken by a light microscope, followed by a quantitative measure via a high sensitivity flow cytometer (BD FACSAria^™^ III) for live/dead cell discrimination. No significant changes in cell viability were observed for cells treated with less or equal to 150 μM of deoxycholic acids

For both HeLa (Graph A) and HuCCT1 (Graph B) cells, a general decrease in the total sia level was observed as the amount of deoxycholic acid increased. The average total sia level for HeLa cells were 0.0062 mM/mg (0 μM), 0.0037 mM/mg (75 μM) and 0.0027 mM/mg (150 μM). The average total sia level for HuCCT1 cells were 0.0046 mM/mg (0 μM), 0.0027 mM/mg (75 μM) and 0.0024 mM/mg (150 μM). A repeated measure ANOVA determined that the mean sia level for both HeLa (F = 31.110 p < 0.0005) and HuCCT1 (F = 16.787, p < 0.005) cell line differed statistically significantly as the deoxycholic acid concentration increased. Therefore, as represented in figure 3, it is concluded that the average amount of sia decreased for 75 μM and 150 μM trials were statistically significantly different to the control group (p < 0.05).

**Figure 3.**
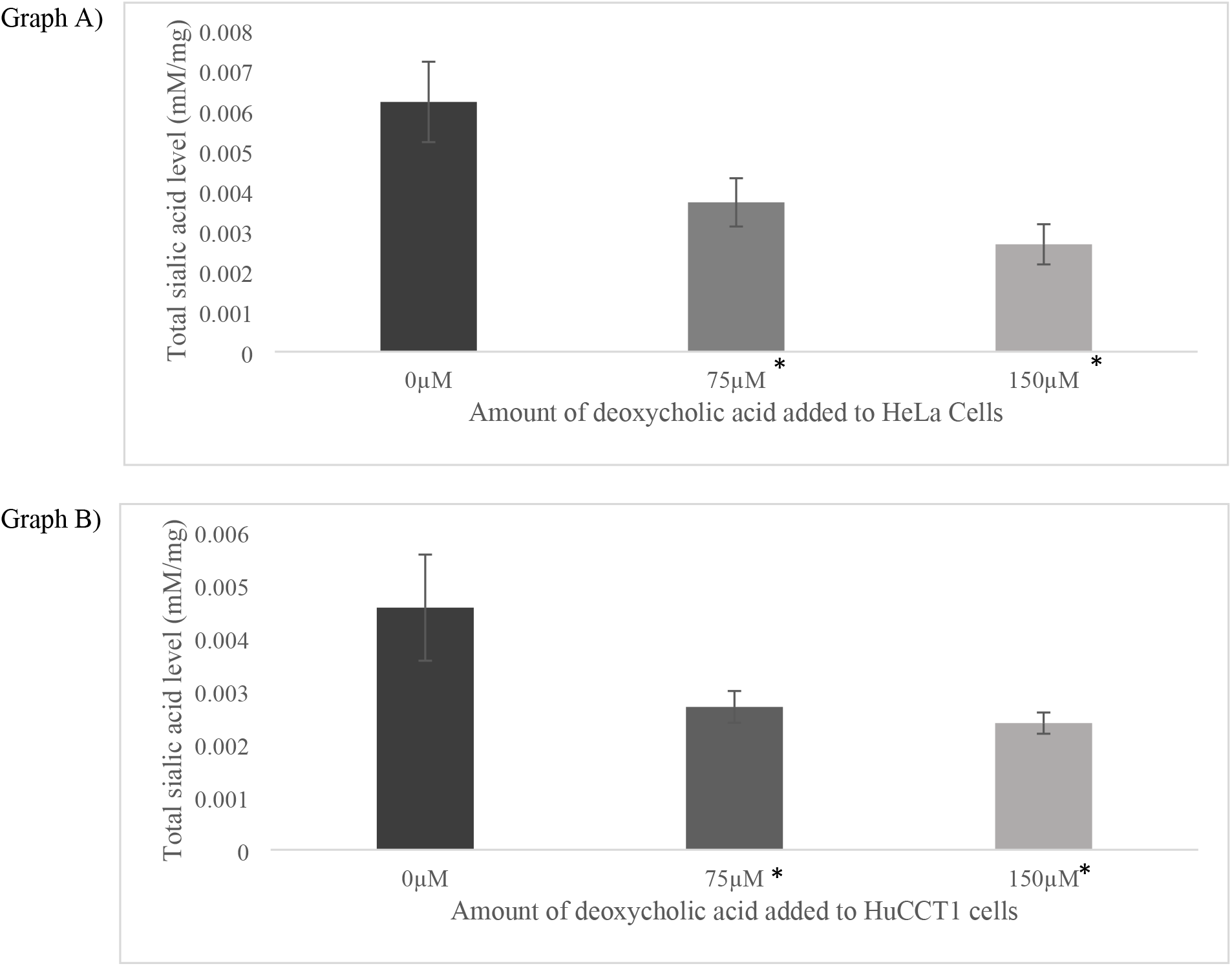
The changes in the total sialic acid level according to various concentration of deoxycholic acid (0 μM, 75 μM and 150 μM) added for HeLa (Graph A) and HuCCT1 (Graph B) cells. The values are mean ± SD, n=5. The asterisks indicate the total sialic acid level for 75 μM and 150 μM were statistically significantly different to the control Neu5Ac level: *P < 0.05.

Figure 3 suggests that as the level of deoxycholic acid increases, the level of sia decreases. This trend suggests that deoxycholic acid, a known lithocholic analogue sialyltransferase inhibitor, has successfully inhibited sialylation in both HeLa and HuCCT1 cells where higher concentration of deoxycholic acid was linked with the extent of sialylation. Therefore, the proposed HPLC method seems to reliably measure the extent of sialylation where it could be used to evaluate potent sialyltransferase inhibitors.

## Discussion and conclusions

This research aimed to develop a method to detect and quantify sias in cancer cells. This method may find a use in evaluating the effectiveness of potent hypersialylation inhibitors. Sias were released and purified through an isolation technique with ethanol precipitation and acid hydrolysis. The samples were then detected and quantified via reverse phase HPLC with TIP buffer solution as the ion-pairing reagent. The proposed method resulted in Neu5Ac eluted with the retention time of 6.344 minutes with a flow rate of 0.4 mL/min. The proposed method was validated appropriately according to the AOAC guideline (2013). This work demonstrates that the proposed method is not only relatively simple but also cost and timely effective compared to pre-existing methods to successfully determine both free and protein-bound Neu5Ac in a complex cancer cell matrix. Furthermore, as suggested by figure 3, the proposed method seems to be applicable in reliably measuring the extent of sialylation in malignant cells to evaluate the effectiveness of potent sialyltransferase inhibitors.

## Availability of data and material

The datasets generated during and/or analysed during the current study are available from the corresponding author on reasonable request.

## Code availability

HPLC system controlled by Empower Chromatography Data software using Ethernet communications.

## Ethics approval

Not applicable.

## Consent to participate

Not applicable

## Consent for publication

Not applicable

## Author contribution

All authors contributed to the study conception and design. Material preparation, data collection and analysis were performed by Hyo Jeong Kim, Stephan Levonis and Stephanie Schweiker. The manuscript was drafted by Hyo Jeong Kim and all authors commented on previous versions of the manuscript. All authors read and approved the final manuscript.

## Acknowledgements

N/A

